# First chromosome scale genome of *Acrocomia aculeata*

**DOI:** 10.1101/2025.05.12.653347

**Authors:** Matheus Scaketti, Caroline Bertocco Garcia, João Victor da Silva Rabelo Araujo, Igor Araujo Santos de Carvalho, Kauanne Karolline Moreno Martins, Diego Mauricio Riaño Pachón, Durval Dourado Neto, Carlos Alberto Labate, Renato Vicentini dos Santos, José Baldin Pinheiro, Suelen Alves Vianna, Carlos Colombo, Maria Imaculada Zucchi

## Abstract

The Arecaceae family comprises economically, and ecologically significant palm species widely distributed across tropical and subtropical regions. Among them, *Acrocomia aculeata*, commonly known as Macauba, has gained attention due to its high oil yield, environmental adaptability, and potential applications in sustainable agriculture and bioenergy production. This study presents the first chromosome scale genome assembly of *A. aculeata* and the first reference genome of the genus, using Oxford Nanopore Technologies sequencing, PacBio HiFi and Hi-C proximity ligation data. The genome covers 1.94 Gbp in the 15 pseudochromosomes, with N50 of 143.43 Mbp, achieving a base quality (QV) of 76.4 and a k-mer completeness of 90.58% of PacBio HiFi data, with k=21. Interspersed repetitive regions represent 77.36% of the total genome, with long terminal repeat (LTRs) accounting for 54.98%. A total of 28.215 protein-coding genes were predicted. The genome was validated with BUSCO, achieving a completeness of 98.9% for predicted proteins. The first chromosome scale genome of Macauba provide new insights and breakthroughs for studies of conservation genomics and plant breeding programs.

## Background & Summary

*Acrocomia aculeata*, commonly known as Macauba, macaw palm, or coyol, is a palm species in the order Arecales order within angiosperm. Native from Mexico to northern Argentina and the Caribbean, *A. aculeata* is highly adaptable to various environments^1^. In Brazil, it thrives in the Cerrado, Atlantic Forest, and Pantanal phytogeographic domains, particularly in pasturelands and degraded areas, where it plays a role in ecological restoration^2^. This adaptability position it as a strategic species for sustainable agriculture and reforestation programs throughout Latin America^3^.

One of the most valuable traits of Macauba is its high oil yield, comparable to *Elaeis guineensis*. The mesocarp oil content can range from 20% to 75% and is rich in monounsaturated fatty acids, predominantly oleic acid (∼49.3%), while the kernel oil ranges from 50% to 60% and contains high levels of lauric acid (∼45.1%)^4^. This composition allows for diverse applications in biodiesel production, food products, and cosmetics, strengthening Macauba’s role in sustainable economic development^5^. As a consequence, it has received attention as an alternative and sustainable source of biofuels and high value bioproducts, presenting potential benefits for reducing environmental impact and promoting renewable energy^6^.

In this context, breeding initiatives for Macauba are gaining strength due to the species’ potential. Until now, there are some studies^7–10^ showing that Macauba can be used in a selective breeding strategy. Although all these efforts, genomics resources for palm species remain scarce, with only 21 published genomes within Arecales, for 9 different species^11^, with just *Elaeis oleifera* being native to Brazil^1^.

To help fill this gap, we report the first chromosome scale of one individual of the important socio-economical species *A. aculeata* using Oxford Nanopore Technologies (ONT), Pacific Biosciences HiFi (PacBio) and Hi-C technology. This is also the first genome of the genus and can provide new insights for the studies of the genetic structure and the evolution of the species, allowing new breakthroughs for both conservation genomics and plant breeding programs.

## Methods

### Sample collection, library construction and sequencing

A young sapling of *A. aculeata was* collected from IAC (Agronomic Institute of Campinas). Nearly 9.5g of young leaves (30 leaves) of an approximated 24-month-old palm tree were freshly collected after a 48-hour dark treatment, immediately frozen in liquid nitrogen, and stored at –80 °C until further processing.

High-molecular-weight (HMW) DNA was extracted using a protocol based on Jones et al.^12^ and the yielded HMW DNA was used for all ONT libraries. HMW DNA samples were purified with AMPure XP beads (Beckman Coulter Inc., CA, United States), and DNA fragments ranging from 10 to 25 Kbp were selectively depleted using the SRE kits from PacBio and ONT. Quality control was performed using a NanoDrop® ND-1000 spectrophotometer (Thermo Fisher Scientific, MA, USA), and the A260/280 and A260/230 ratios were assessed to determine DNA purity. Genomic DNA concentration was confirmed using a Qubit® 3.0 fluorometer (Invitrogen, CA, USA) with the dsDNA BR Assay Kit (Thermo Fisher Scientific, MA, USA, Q32850). The integrity and size of high-molecular-weight (HMW) DNA fragments were evaluated using the 4200 TapeStation system (Agilent Technologies, Santa Clara, CA, USA).

ONT long-read sequencing was performed using the SQK-LSK114 kit and three FLO-PRO114M flow cells on the P2-Solo platform, yielding 87.7 Gbp of raw reads, with a mean read size of 5.91 Kbp and read N50 of 10.48 Kbp. All ONT data were basecalled using the dna_r10.4.1_e8.2_400bps_sup@v4.2.0 model. Reads shorter than 5 Kbp or with a Q score below 7 were filtered out. To improve read quality, an error correction step was performed using Haplotype-aware ERRor cOrrection (HERRO)^13^ tool, resulting in 45 Gbp of corrected reads. To simulate an Ultra-long ONT sequencing, only reads longer than 30 Kbp were retained for further analyses.

For PacBio sequencing and High-throughput Chromosome conformation capture (Hi-C), leaf tissue samples were sent to the Arizona Genomics Institute, where nuclei isolation, DNA and HMW DNA extraction were done. Before library preparation, fragments sizes between 10 and 20 kb were enriched using BluePippin, later, two SMRTbell libraries were constructed with the SMRTbell Prep Kit 3.0 (Pacific Biosciences, CA, USA). Sequencing was performed on the REVIO platform at AGI, generating 234.64 Gbp of HiFi reads with a mean read size of 17 Kbp and read N50 of 18 Kbp, to ensure high-quality genome assembly. The Hi-C library was performed using the High Coverage Hi-C kit (Arima Genomics), which proprietary restriction enzymes cuts at ^GATC, G^ANTC, C^TNAG, and T^TAA sites. Hi-C library sequencing was conducted by Arima Genomics on Illumina NovaSeq X Plus platform, generating 69.6 Gbp of raw data of 2x150bp (461.4 million reads).

### *De novo* genome assembly and chromosome level scaffolding

Genome features were estimated with GenomeScope2^14^, by computing a k-mer spectra using Meryl^15^, and on the PacBio HiFi reads. *A. aculeata* is a diploid species (2n = 30) with an estimated genome size of 2,84 Gbp, based on the average 2C value of 5,81 pg^16^. However, GenomeScope2 results estimates a monoploid genome size of 1.86 Gbp (Supplementary Figure S1), 38.4% of non-repetitive fraction of the genome, and a heterozygosity among the two haplophases of the genome 0.617%. Monge-Castro et al. and Morales-Marroquín et al. shows that Macauba populations presents high levels of population genetic structure, including cases of low gene flow among geographically distant populations, we hypothesize that the repeat content of these populations may vary, and thus, presenting different estimations of the genome sizes of Macauba^17,18^. Considering this, and methodological limitation of both cytometric and k-mer-based approaches, we adopted the GenomeScope2 monoploid estimate as reference for the following analyses, even though this method does not provide an exact measure of genome size in repeat-rich, heterozygous plants genomes.

A draft diploid genome assembly was generated using Hifiasm^19^ v0.24.0-r702 integrating all 30+ Kbp ONT, PacBio HiFi and Hi-C data, applying default parameters. This assembly produced 683 primary contigs with a total length of 2.04 Gbp and an N50 of 141.4 Mbp. Haplotigs from the primary assembly were subsequently removed using purge_dups^20^ v1.2.5, yielding a final draft assembly of 23 contigs with a total length of 1.96 Gbp.

Scaffolding was performed using the purged draft genome and the Hi-C data using the Juicer v1.6 pipeline^21^ with assembly mode, following the instructions provided by Arima Genomics^22^ for the high-coverage kit. Re-scaffolding and interactive Hi-C contact map were generated with YaHS^23^ v1.2.2. Manual curation of contigs and scaffolds was conducted using Juicebox^24^ v2.17, resulting in 27 curated contigs organized into 15 pseudochromosomes, containing 7 gaps.

### Gene prediction and Functional annotation

De novo repeat families were identified and classified using RepeatModeler^25^ v2.0.6 with the -LTRStruct option, resulting in the construction of a custom repeat library. Subsequently, the Viridiplantae dfam database was retrieved and combined with the *de novo* library for masking using RepeatMasker^26^ v4.1.8. We identified 1.51 Gbp of interspersed repeat regions, comprising 77.36% of the genome (Table 1). Among these regions, LTR elements represent the majority, composed of 54.98%, where 35.82% being Ty1/Copia and 14.91%, Gypsy/DIRS1.

**Table 1.**
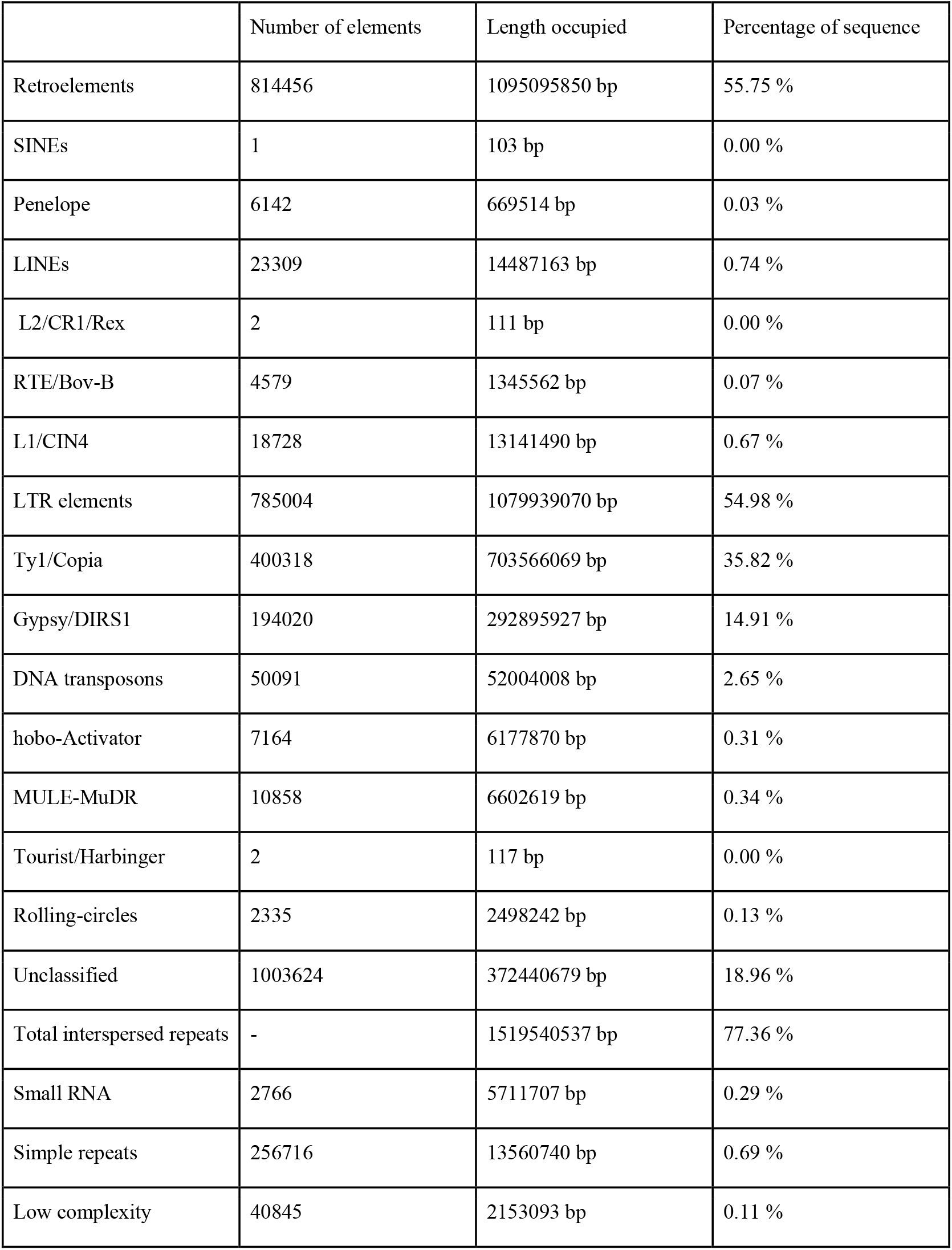
Annotation of repeat content for *Acrocomia aculeata*.

|  | Number of elements | Length occupied | Percentage of sequence |
| --- | --- | --- | --- |
| Retroelements | 814456 | 1095095850 bp | 55.75 % |
| SINEs | 1 | 103 bp | 0.00 % |
| Penelope | 6142 | 669514 bp | 0.03 % |
| LINEs | 23309 | 14487163 bp | 0.74 % |
| L2/CR1/Rex | 2 | 111 bp | 0.00 % |
| RTE/Bov-B | 4579 | 1345562 bp | 0.07 % |
| L1/CIN4 | 18728 | 13141490 bp | 0.67 % |
| LTR elements | 785004 | 1079939070 bp | 54.98 % |
| Ty1/Copia | 400318 | 703566069 bp | 35.82 % |
| Gypsy/DIRS1 | 194020 | 292895927 bp | 14.91 % |
| DNA transposons | 50091 | 52004008 bp | 2.65 % |
| hobo-Activator | 7164 | 6177870 bp | 0.31 % |
| MULE-MuDR | 10858 | 6602619 bp | 0.34 % |
| Tourist/Harbinger | 2 | 117 bp | 0.00 % |
| Rolling-circles | 2335 | 2498242 bp | 0.13 % |
| Unclassified | 1003624 | 372440679 bp | 18.96 % |
| Total interspersed repeats | - | 1519540537 bp | 77.36 % |
| Small RNA | 2766 | 5711707 bp | 0.29 % |
| Simple repeats | 256716 | 13560740 bp | 0.69 % |
| Low complexity | 40845 | 2153093 bp | 0.11 % |

To validate the accuracy of LTR retrotransposon classification, intact LTR elements were independently identified using LTRharvest followed by LTR_retriever on the final genome assembly. The genomic coordinates of LTR/Gypsy/Copia elements annotated by RepeatMasker were then intersected with the structurally defined LTRs using BEDTools v2.30.0. This comparison revealed that 77.48% of the LTR elements annotated by RepeatMasker showed direct structural support, confirming that the high abundance of LTR retrotransposons in the genome is supported not only by homology-based classification but also by independent structure-based evidence.

Although no repeat families remained unclassified at the superfamily level (i.e., all families could be assigned to known Viridiplantae/Dfam TE classes), most of these families represent lineage-specific variants identified de novo in the macauba genome. Therefore, the repeatome expansion observed here reflects extensive species-specific amplification of known TE lineages rather than the origin of entirely new deep-level TE families.

Using the chromosome level assembly, gene prediction was performed incorporating RNA-seq datasets from 21 accessions, comprising different tissues of the palm tree (Table 2) - including fruit pulp, male and female flowers, roots, and leaves (NCBI BioProject PRJNA489676^27,28^). Each dataset was aligned to the 15 pseudochromosomes using HISAT2^29^ v2.2.1 with --rna-strandness RF and --dta, and the resulting alignment files were merged with samtools^30^ v1.21.

**Table 2.** Read mappability across currently available NCBI *A. aculeata*’s RNA-seq datasets.

| Accession number | Sample part | Base pairs | Read number | Mappability (%) |
| --- | --- | --- | --- | --- |
| SRR7822563<sup>31</sup> | Leaf | 4,2 Gbp | 20938255 | 92,47 |
| SRR7822564<sup>32</sup> | Leaf Sheath | 5,9 Gbp | 29394232 | 91,79 |
| SRR7822565<sup>33</sup> | Female Flower | 15,7 Gbp | 78987778 | 81,24 |
| SRR7822566<sup>34</sup> | Female Flower | 5,5 Gbp | 27824979 | 82,24 |
| SRR7822567<sup>35</sup> | Leaf | 4,8 Gbp | 24284522 | 92,39 |
| SRR7822568<sup>36</sup> | Leaf | 4,6 Gbp | 23053873 | 92,54 |
| SRR7822569<sup>37</sup> | Bulb | 4,1 Gbp | 20773350 | 91,67 |
| SRR7822570<sup>38</sup> | Bulb | 5,1 Gbp | 25893211 | 92,60 |
| SRR7822571<sup>39</sup> | Bulb | 4,6 Gbp | 23313242 | 91,47 |
| SRR7822572<sup>40</sup> | Female Flower | 5,7 Gbp | 28635910 | 54,10 |
| SRR7822573<sup>41</sup> | Root | 4,9 Gbp | 24696967 | 83,62 |
| SRR7822574<sup>42</sup> | Root | 4,3 Gbp | 21595079 | 84,03 |
| SRR7822575<sup>43</sup> | Root | 4,7 Gbp | 23561512 | 86,58 |
| SRR7822576<sup>44</sup> | Leaf Sheath | 5,5 Gbp | 27501990 | 93,17 |
| SRR7822577<sup>45</sup> | Leaf Sheath | 5,1 Gbp | 25450315 | 92,81 |
| SRR7822578<sup>46</sup> | Male Flower | 4,5 Gbp | 22564627 | 76,61 |
| SRR7822579<sup>47</sup> | Male Flower | 5 Gbp | 24850587 | 72,59 |
| SRR7822580<sup>48</sup> | Male Flower | 4,5 Gbp | 22487082 | 85,94 |
| SRR7822581<sup>49</sup> | Ripening fruit | 6,4 Gbp | 31804017 | 32,90 |
| SRR7822582<sup>50</sup> | Ripening fruit | 5,9 Gbp | 29007834 | 47,66 |
| SRR7822583<sup>51</sup> | Ripening fruit | 6,5 Gbp | 31986486 | 36,21 |

After processing each RNA-seq data, we analyzed the mappability percentage for each tissue. The percentage of reads aligned to the assembled genome varied from 36.21% to 92.47%, with an average of 78.79%. Leaf tissues, female flowers and bulb were the plant parts that got the highest mapped reads, indicating a high level of completeness and structural accuracy in genic regions, allowing effective transcriptome-supported annotation.

The quality and coverage of the aligned reads for each transcript were assessed using QualiMap52 v2.2.2, to avoid misleading extrinsic evidence from RNA-seq quality, we filtered out datasets with read mappability lower than 60% (SRR7822572, SRR7822581, SRR7822582 and SRR7822583). Lower mappability in some RNA-seq libraries is likely due to reduced RNA quality, particularly in samples derived from fruit tissues, which are rich in oils and secondary metabolites that negatively affect RNA extraction and downstream sequencing. NCBI RefSeq protein sequences for Elaeis guineensis were downloaded (48.771 sequences).

Both RNA-seq and protein data were used as extrinsic evidence for gene prediction using BRAKER3^53–59^ v3.0.8 with --softmasking and --gff3 parameters. The resulting gene prediction was then filtered using AGAT^60^ v1.0.0 to retain only genes with transcript support and remove conflicting features, maintaining only the longest isoform for each predicted gene and reducing the redundancy, resulting in a total of 28.215 predicted genes.

To functionally annotate the predicted proteins, we utilized EggNOG-mapper^61^ v2.1.8, PANNZER2^62^ and Automated Assignment of Human Readable Descriptions (AHRD) tool^63^ v3.3.3 function descriptions, providing functional insights into the predicted gene set. EggNOG annotation was unable to identify 2.382 isoforms and AHRD defined 995 isoforms as Unknown proteins. Considering all three methods, all predicted proteins were functionally annotated by at least one source.

### Identification of non-coding RNAs

We employed a combination of curated tools optimized for different RNA classes. Firstly, we used tRNAscan-SE^64^ v1.4 with default parameters to locate transfer RNA (tRNA) genes. Ribosomal RNAs (rRNA) and other non-coding RNAs were predicted using the RFAM database with Infernal’s cmscan^65^ v1.1.5 command, filtering out predictions with E-value greater than 0.05. A total of 5761 rRNA, 1679 tRNAs, 932 snoRNAs, 162 miRNAs and 138 snRNAs were identified.

### Data overview

The final assembly is highly contiguous, with a N50 of 143.43 Mbp and a scaffold assembly of all 15 pseudochromosomes covering a total of 1.94 Gbp, with lengths ranging from 79.45 to 175.71 Mbp (Table 3). The Telomere-to-Telomere toolkit^71^ v1.2.5 (quarTeT) and Telomere Identification ToolKit^72^ v0.2.63 (TIDK) were employed to identify and create the plot to visualize telomeric repeats across all pseudochromosomes, resulting in the sequence AAACCCT. We assembled nine pseudochromosomes with telomeric repeats on both ends, and the remaining six with a telomeric repeat at only one end.

**Table 3.** Summary of genome metrics for *Acrocomia aculeata*.

| Assembly | Genome length | Contigs number | N50 | Assembly BUSCO | Proteins BUSCO | QV | K-mer completeness | False duplication |
| --- | --- | --- | --- | --- | --- | --- | --- | --- |
| contig assembly | 1.96 Gbp | 27 | 143.43 Mbp | 99.4% | - | 75.66 | 90.69% | 1.72% |
| scaffold assembly | 1.94 Gbp | 15 | 143.43 Mbp | 99.4% | 98.9% | 76.40 | 90.58% | 1.71% |

Pseudochromosome structure and genomic features were visualized using a CIRCOS plot, displaying gene density, Gypsy and Copia elements densities, GC content and pseudochromosome collinearity. Collinearity analyses were performed with MCScanX^68^ v1.0.0 (Figure 1).

**Figure 1:**
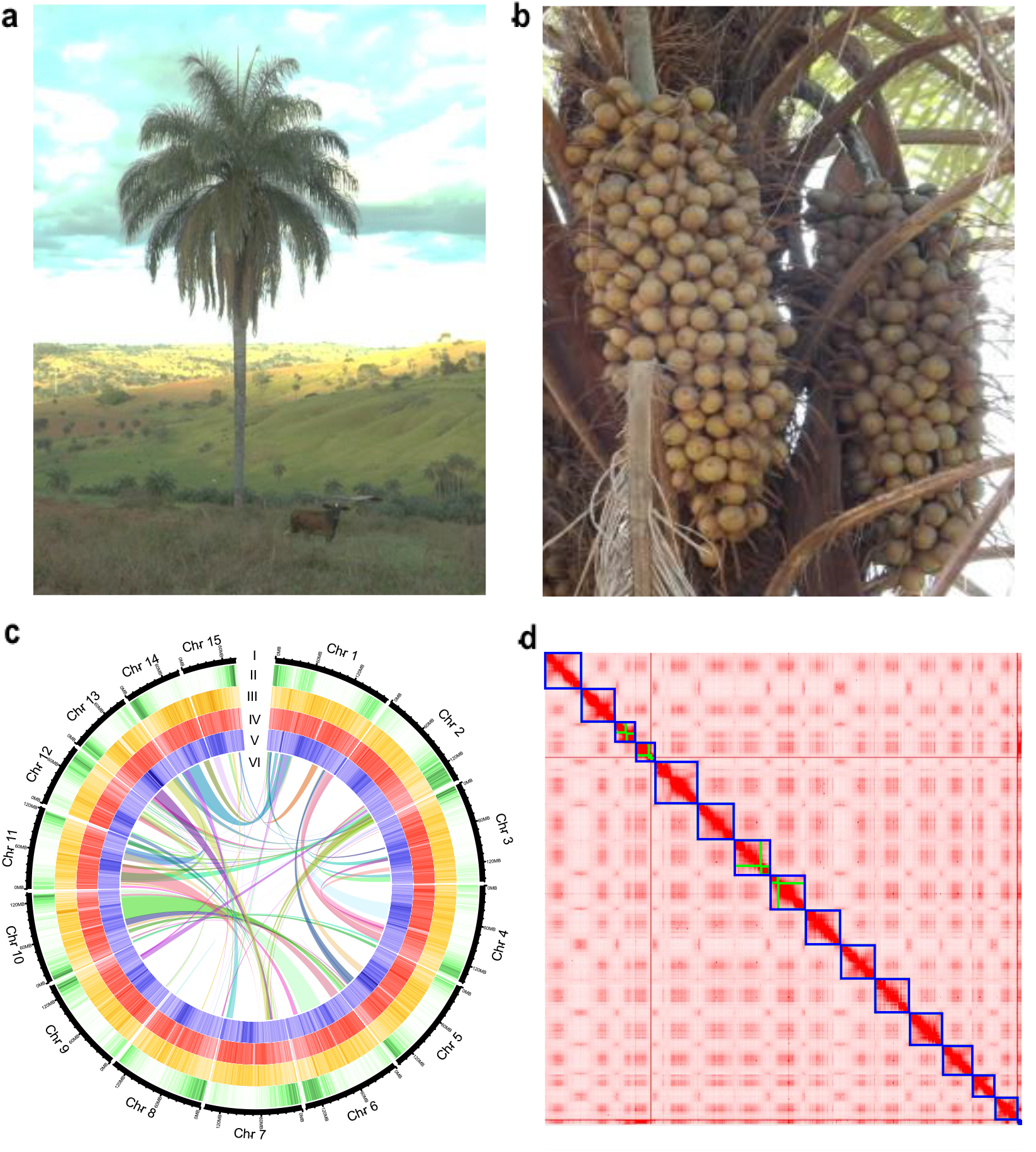
Morphology and genomic features of *Acrocomia aculeata*. **(A)** Adult individual, located at Carmo do Paranaiba, Minas Gerais, in frontal view, showing a solitary stipe and pinnately compound leaves. **(B)** Infructescence bearing globose drupes with a yellow epicarp. **(C)** Circos plot of the *A. aculeata* genome; tracks represent (I) chromosomes, (II) gene density, (III) LTR/Gypsy element density, (IV) LTR/Copia element density, (V) GC content, and (VI) chromosome collinearity blocks (self vs. self). **(D)** Hi-C contact map illustrating chromatin interactions across the genome. Photographs a) and b) taken by Suellen Alves Vianna.

### Technical Validation

To quantify the impact of the HERRO correction step, we compared raw and corrected ONT reads using kmer-based metrics (Merqury) and alignment statistics. Raw ONT reads showed a mean alignment identity of 95.3% and a median of 98.4%, with 61.6% of bases above Q20 and a total yield of 95.3 Gb, as reported by NanoPlot. After HERRO correction, the k-mer completeness increased from 87.25% to 91.35%, indicating improved recovery of genomic content.

Gene space completeness of the genome assembly was assessed using BUSCO^66^ v5.7 with the *liliopsida_odb12* lineage, contiguity metrics were assessed with QUAST^67^ v5.3.0 (Supplementary Figure S2). Base-level quality (QV), k-mer completeness and false duplication percentage were estimated with Merqury^15^ v1.3 using PacBio HiFi data, considering k=21.

Comparing the gene amount found is to other palm species such as the Oil palm, Coconut and Date palm, which has gene counts of 28.077, 28.000 and 29.239 predicted genes, respectively (Table 4). Gene structural features are also similar, with *A. aculeata* showing an average of 7 exons per mRNA and a mean gene length of 7.697 bp, values intermediate between *E. guineensis* and *P. dactylifera* and slightly lower than those observed in *C. nucifera*. Mean CDS length (1,314 bp) and exon length (228 bp) in *A. aculeata* fall within the narrow range observed among the other palms, supporting consistency in coding sequence architecture. Together, these comparisons indicate that the gene prediction and annotation of *A. aculeata* are structurally coherent and comparable to well-annotated reference palm genomes.

**Table 4.** Annotation summary of palm species.

|  | *A. aculeata* | *E. guineensis* | *C. nucifera* | *P. dactylifera* |
| --- | --- | --- | --- | --- |
| Number of genes | 28.215 | 28.077 | 28.000 | 29.239 |
| Number of mrnas | 34.516 | 48.771 | 28.000 | 48.801 |
| Mean mrnas per gene | 1.2 | 1.7 | 1.0 | 1.7 |
| Mean cds per mrna | 1.0 | 1.0 | 1.0 | 1.0 |
| Mean exons per mrna | 5.7 | 7.2 | 4.7 | 7.0 |
| Mean exons per cds | 5.7 | 6.5 | 4.7 | 6.4 |
| Mean gene length | 7.697 | 9.325 | 6.635 | 8.123 |
| Mean mrna length | 8.222 | 12.179 | 6.635 | 10.103 |
| Mean cds length | 1.314 | 1.419 | 1.155 | 1.432 |
| Mean exon length | 228 | 297 | 345 | 305 |
| Longest gene | 438.084 | 266.372 | 187.278 | 492.108 |

Corrected ONT reads and raw HiFi reads were aligned to the final genome assembly, resulting in 91.3% and 93.3% of aligned bases and mean sequence identities of 95.3% and 96.8%, respectively. The BUSCO analysis showed that our assembly has 79.3% and 20.1% of complete and duplicated orthologues genes, with only 0.3% and 0.3% fragmented or missing. Alongside that, a low false duplication of 1.71%, K-mer completeness of 90.58% and QV greater than 75, demonstrates a high-quality assembly. Gene prediction with both RNA-seq and *E. guineensis*’ proteins, has a BUSCO completeness of 98.9%, with 79.6% single copy and 19.3% duplicated.

To access the genome assembly quality on repetitive regions, we computed the LTR assembly index (LAI), resulting in an average raw score of 25.11, being considered an “gold standard” reference genome^69^. This result shows that our genome is comparable to the recent reference genome of *E. guineensis* and *E. oleifera*, with LAI scores of 15.33 and 18.93^70^.

In addition, we compared the 15 pseudochromosomes of *A. aculeata* against the genomes of four palm species (Figure 2): *Elaeis guineensis, E. oleifera*^70^, *Cocos nucifera*^73^ and *Phoenix dactylifera*^74^, using D-Genies web service^75^. These results show higher levels of collinearity between *A. aculeata* and *E. guineensis, E. oleifera* and *C. nucifera* than with *P. dactylifera* which is expected, because these palm species are from the same subfamily Cocosaea. Finally, for *P. dactylifera* which are from Coryphoideae, there are more differences between chromosomes, containing fragmented collinearity regions^76^.

**Figure 2:**
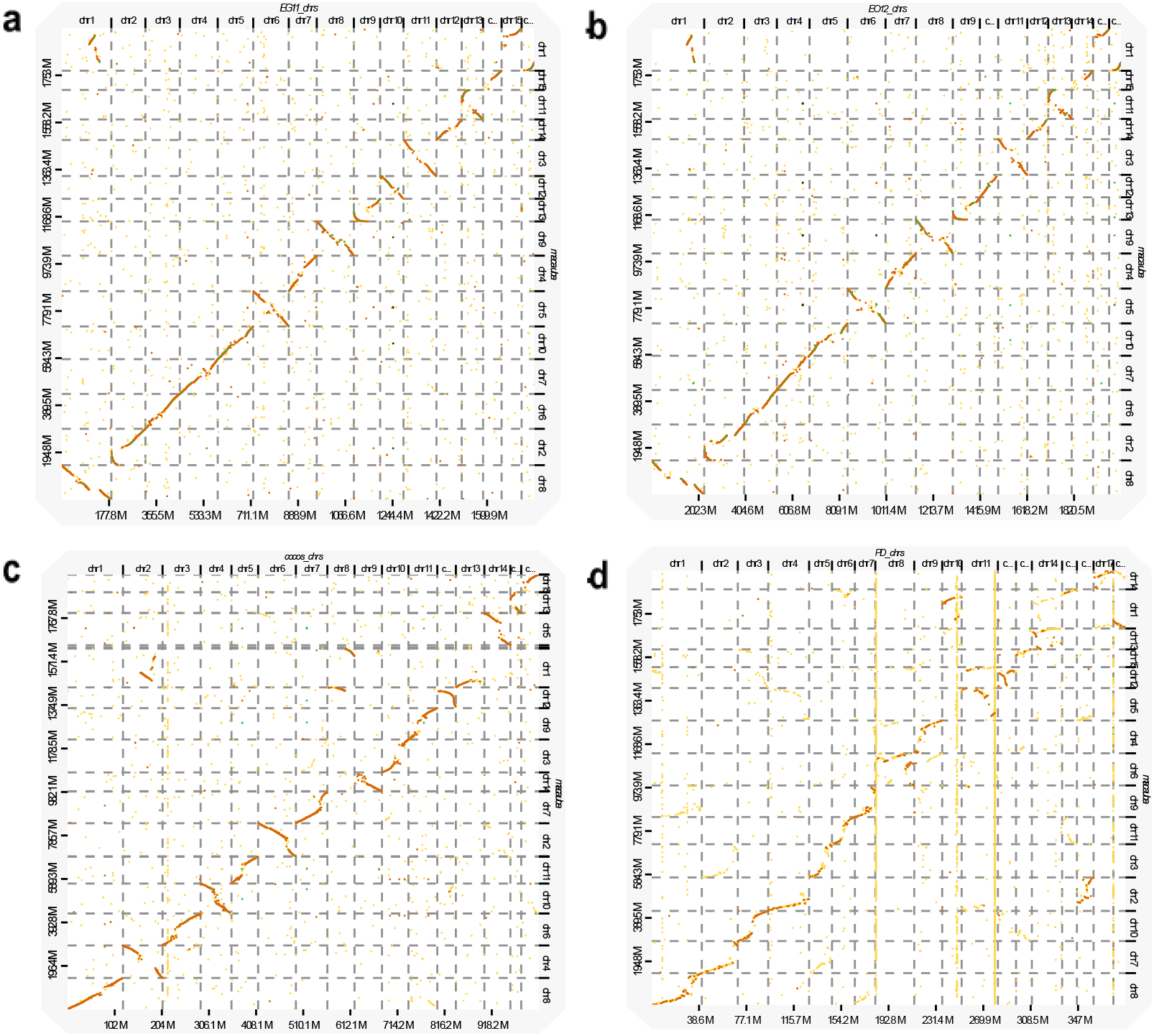
Base-by-base comparison between *Acrocomia aculeata* and other palm species. (a) *Elaeis guineensis* (African oil palm; EG11; PRJNA192219^77^). (b) *Elaeis oleifera* (American oil palm; EO12.1; PRJNA183707^78^).(c) *Cocos nucifera* (Coconut palm; Cnuc_CPCRI_v2; PRJNA413280^79^). (d) *Phoenix dactylifera* (Date palm; Barhee BC4; PRJNA322046^80^).

It is possible to observe some chromosome rearrangements (highlighted in red). *E. guineensis* and *A. aculeata* shows inverted regions between chromosome 2 and 8 and a spliced region shared with chromosome 1, also having inverted collinearity between pseudochromosomes 3, 5, 9 and 11 and chromosomes 11, 6, 8 and 13. *Elaeis oleifera* exhibits a collinearity pattern largely congruent with *E. guineensis*, but with distinct differences in each chromosome, such as chromosomes 1 and 2. *C. nucifera* presents less similar regions than *E. guineensis* and *E. oleifera*, with more rearrangements, found at pseudochromosomes 1, 2, 4, 5, 10 and 13. Pseudochromosome 1 also shows fragmentation between different *C. nucifera* chromosomes 2 and 8. Finally, the comparison between *P. dactylifera* reinforces the phylogenetic distance between the species, with high fragmented regions such those found at pseudochromosomes 1, 2, 4, 5, 6, 12, 13 and 14.

To further validate the genome assembly, we conducted comparative phylogenetic and gene synteny analyses (Figure 3) with closely related palm species, *Elaeis guineensis* (Eg), *Cocos nucifera* (Cn) and *Phoenix dactylifera* (Pd). We used GENESPACE^81^ v1.2.3 to access the phylogenomic inference based on genome-wide orthologous genes, our results recovered a well-supported topology in which *A. aculeata* clusters as a close species to *E. guineensis* and with *C. nucifera* forming a closely related lineage. *P. dactylifera* placed as the outgroup. This topology (Supplemental Figure S3) is fully consistent with established phylogenetic relationships within Arecaceae and supports the evolutionary placement of *A. aculeata* among oil-producing palms^82^.

**Figure 3:**
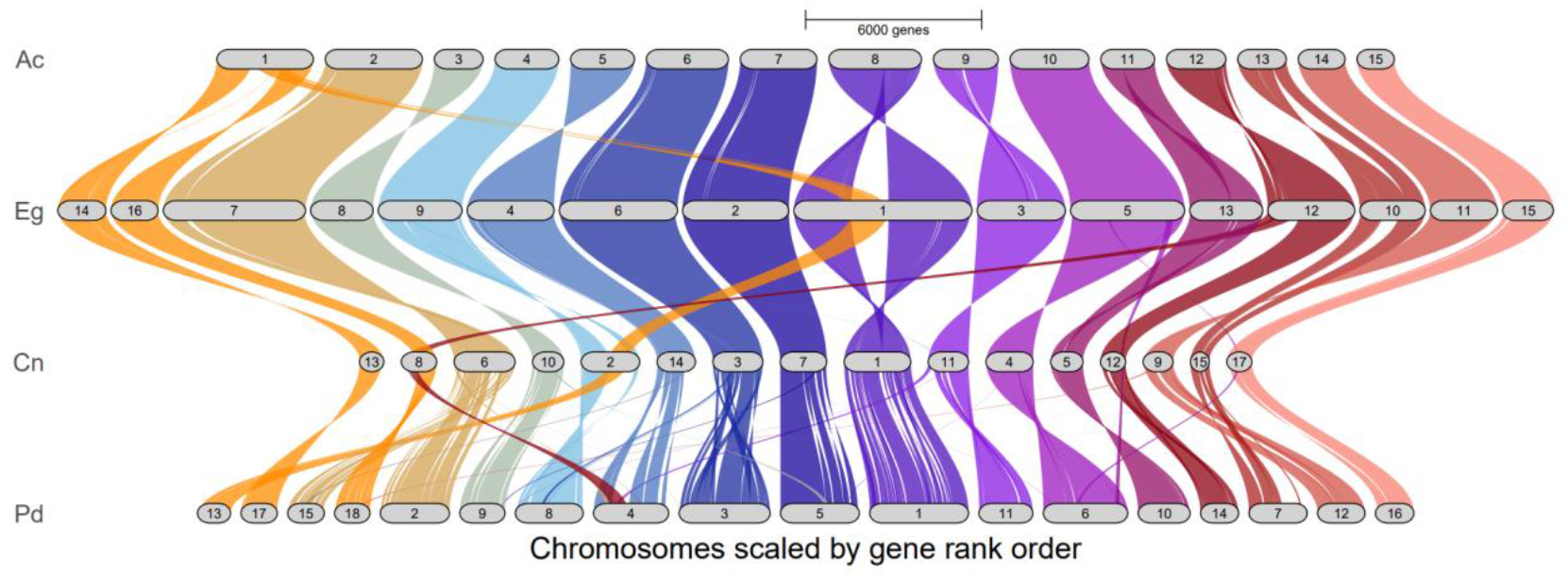
Gene orthology-based synteny between *A. aculeata* and other palm species. (Ac) *Acrocomia aculeata* (This paper). (Eg) *Elaeis guineensis* (African oil palm; EG11; PRJNA192219). (Cn) *Cocos nucifera* (Coconut palm; Cnuc_CPCRI_v2; PRJNA413280). (Pd) *Phoenix dactylifera* (Date palm; Barhee BC4; PRJNA322046).

Gene synteny analyses revealed a chromosome-scale collinearity between *A. aculeata* and the other palm genomes. Large syntenic blocks were conserved across all comparisons, with chromosomes largely maintaining one-to-one correspondence in gene order and orientation. Although several chromosomal rearrangements and translocations were detected, such as found in chromosome 1 of Macauba being divided into the chromosomes 1, 14 and 16 of *E. guineensis*, these were limited in number and are consistent with lineage-specific structural evolution rather than assembly artifacts. The overall conservation of gene rank order across chromosomes further supports the structural accuracy of the assembly.

## Code availability

No custom code was created for this manuscript; we used publicly available programs with default parameters; different options were described in the Methods.

## Acknowledgements

Acelen Renovaveis supported this work by funding the costs of the PacBio HiFi and Hi-C sequencing data (Project number: FEALQ 104.781). FAPESP 2021/10319-0 and CNPq 313417/2023-7 also supported this work for all ONT sequencing data.

## Author contributions

M.S. and M.I.Z. conceptualized and designed this study and manuscript review. C.B.G., and M.S, HMW DNA extraction, library preparation, Nanopore DNA sequencing and manuscript preparation. C.B.G., I.A. and J.R. CTAB DNA extraction and Illumina library preparation., M.I.Z., D.D.N., C.A.L., J.B.P. funding coordination, C.C. sample collection. M.S. technical validation, bioinformatic analysis and data submission. K.K.M.M., S.A.V., D.M.R.P. and R.V.S. manuscript review.

## Competing interests

The authors declare no competing interests.

## Additional information

## Notes

### Competing Interest Statement

The authors have declared no competing interest.

### Summary of Updates

We added more validations to genome quality, as well as gene synteny and phylogenomic information between other palm species.

