## Supplementary Files for "First chromosome scale genome of *Acrocomia aculeata*"

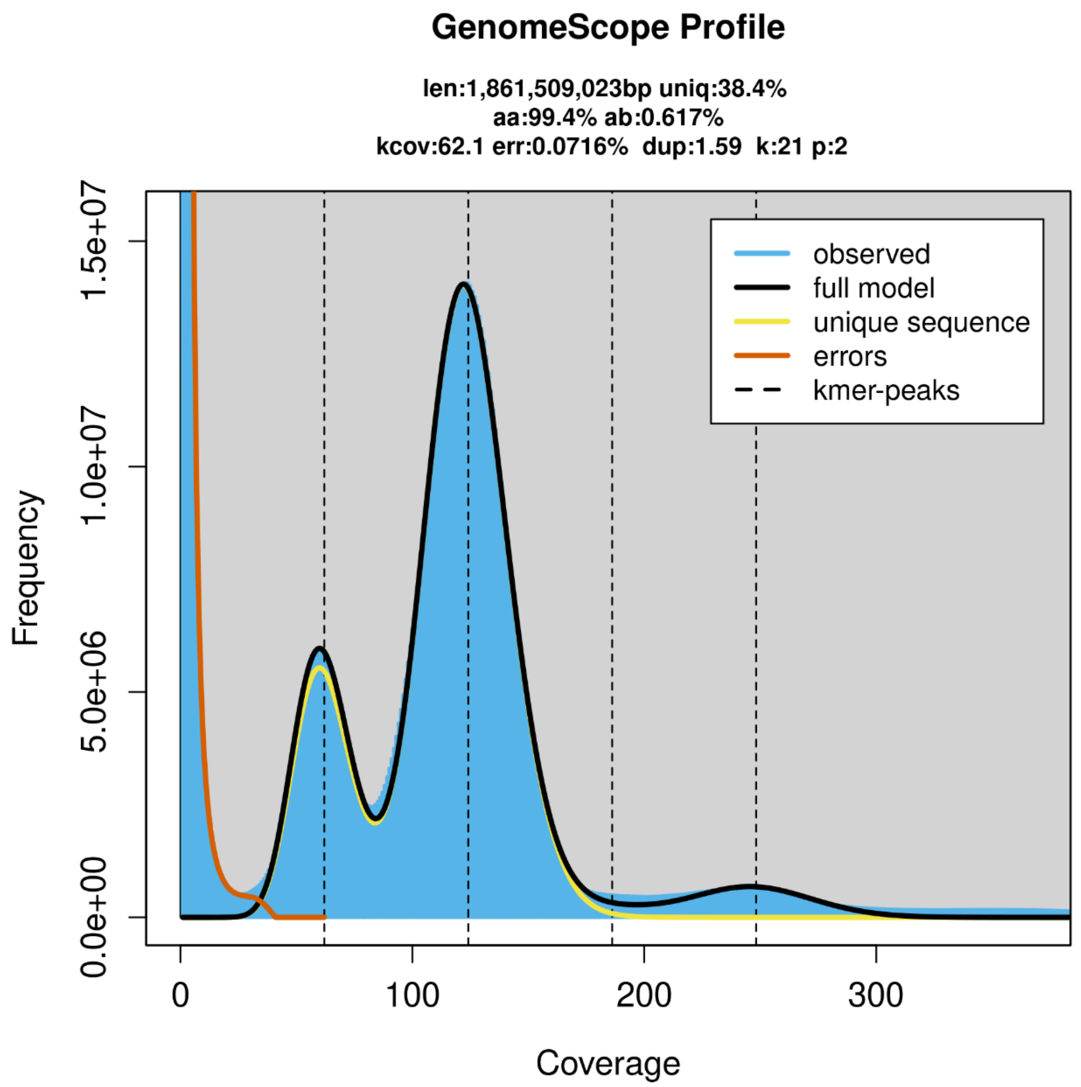


**Figure S1.** Genome characteristics of *Acrocomia aculeata* inferred from k-mer frequency distribution using GenomeScope2. The analysis estimated a monoploid genome size of 1.86 Gbp, with 38.4% unique sequence content and a heterozygosity rate of 0.617%. The model indicates an average k-mer coverage of 62.1×, low sequencing error (0.0716%), and limited duplication (1.59%), supporting the accuracy of the sequencing data and overall genome complexity.

A

| | Contigs Assembly |
|---|---|
| # contigs (>= 0 bp) | 27 |
| # contigs (>= 1000 bp) | 27 |
| # contigs (>= 5000 bp) | 27 |
| # contigs (>= 10000 bp) | 24 |
| # contigs (>= 25000 bp) | 22 |
| # contigs (>= 50000 bp) | 20 |
| Total length (>= 0 bp) | 1964193835 |
| Total length (>= 1000 bp) | 1964193835 |
| Total length (>= 5000 bp) | 1964193835 |
| Total length (>= 10000 bp) | 1964178835 |
| Total length (>= 25000 bp) | 1964148835 |
| Total length (>= 50000 bp) | 1964050296 |
| # contigs | 27 |
| Largest contig | 175714414 |
| Total length | 1964193835 |
| GC (%) | 39.95 |
| N50 | 143438751 |
| N90 | 93622483 |
| auN | 134599248.4 |
| L50 | 7 |
| L90 | 13 |
| # N's per 100 kbp | 0.04 |

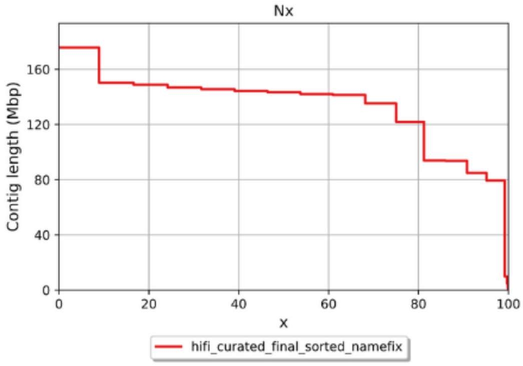


B

| | Scaffold Assembly |
|---|---|
| # contigs (>= 0 bp) | 15 |
| # contigs (>= 1000 bp) | 15 |
| # contigs (>= 5000 bp) | 15 |
| # contigs (>= 10000 bp) | 15 |
| # contigs (>= 25000 bp) | 15 |
| # contigs (>= 50000 bp) | 15 |
| Total length (>= 0 bp) | 1947742739 |
| Total length (>= 1000 bp) | 1947742739 |
| Total length (>= 5000 bp) | 1947742739 |
| Total length (>= 10000 bp) | 1947742739 |
| Total length (>= 25000 bp) | 1947742739 |
| Total length (>= 50000 bp) | 1947742739 |
| # contigs | 15 |
| Largest contig | 175714414 |
| Total length | 1947742739 |
| GC (%) | 39.85 |
| N50 | 143438751 |
| N90 | 93622483 |
| auN | 135673386.9 |
| L50 | 7 |
| L90 | 13 |
| # N's per 100 kbp | 0.04 |

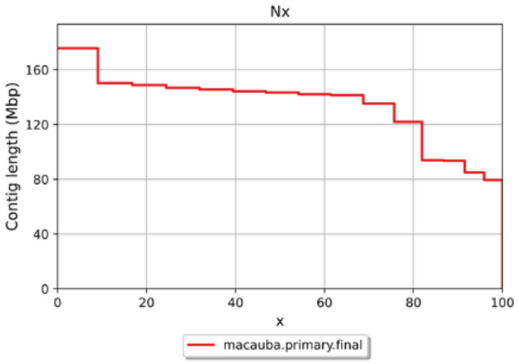


**Figure S2.** Assembly statistics and contiguity metrics for the *Acrocomia aculeata* genome generated with QUAST. (A) Contig-level assembly showing 27 contigs totaling approximately 1.96 Gbp. (B) Scaffold-level assembly organized into 15 pseudochromosomes spanning about 1.94 Gbp. Nx plots depict the cumulative contig and scaffold length distributions, confirming the high contiguity of the chromosome-scale assembly.

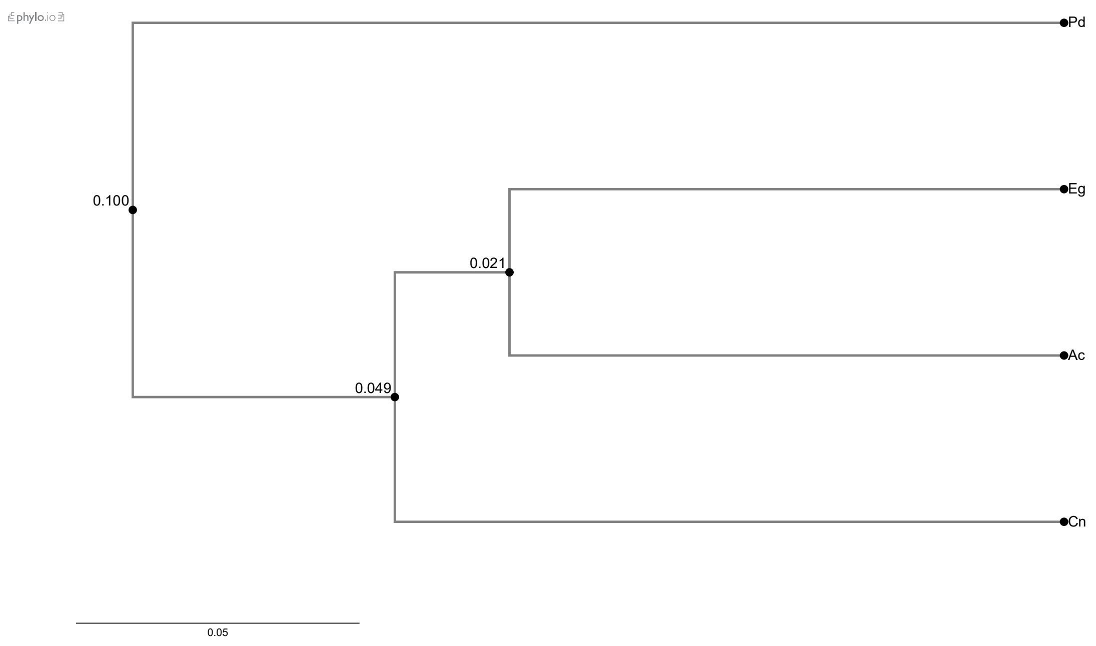


**Figure S3.** Phylogenomic inference based on genome-wide orthologous genes among *Acrocomia aculeata* and related palm species. The topology recovers *A. aculeata* as a lineage closely related to *Elaeis guineensis*, with *Cocos nucifera* forming a neighboring clade, while *Phoenix dactylifera* is positioned as the outgroup. The conserved branching pattern agrees with established evolutionary relationships within Arecaceae and provides independent support for the structural accuracy of the assembled genome.
